## Supplemental Information for "Comparative Interactome Analysis of α-arrestin Families in Human and *Drosophila*"

**Supplementary Methods** ------------------------------------------------------------------------ Page 3

**Supplementary Figures**

**Figure S1.** Fluorescence images showing HEK293 and S2R+ cells stably expressing GFP-tagged α-arrestins -------------------------------------------------------------------------------- Page 10

**Figure S2.** AP/MS data offer high reproducibility and expand the PPIs associated with α-arrestins while also reaffirming known interactions ------------------------------------------ Page 11

**Figure S3.** Protein domains and subcellular localization of α-arrestin interactomes -- Page 13

**Figure S4.** Network of high-confidence PPIs between α-arrestins and individual proteins of its associated protein complexes ----------------------------------------------------------------- Page 15

**Figure S5.** Protein sequence homology of α-arrestins from human and *Drosophila* -- Page 18

**Figure S6.** High-throughput sequencing data are highly reproducible and ATAC-seq reads exhibit a typical pattern of strong enrichment around TSSs of expressed genes ------ Page 19

**Figure S7.** Genomic locations of ACRs and association between chromatin accessibility and transcriptional activity --------------------------------------------------------------------------- Page 20

**Figure S8.** TXNIP might play role in transcriptional regulation independent of known factors --- ---------------------------------------------------------------------------------------------------- Page 22

**Figure S9.** Association between α-arrestin interactomes and human diseases -------- Page 24

**Supplementary Table and legends** ------------------------------------------------------------- Page 25

**References** ----------------------------------------------------------------------------------------------- Page 30

### Supplementary Methods

#### Immunofluorescence imaging of human α-arrestins

Stably α-arrestin-GFP expressing HEK293 cells were cultured in a 12 well-plate with pre-sterilized round glass coverslips in each well. Cells on coverslip were fixed in 4% paraformaldehyde (PFA) (RT15710, Electron Microscopy Sciences, Hatfiled, PA, USA) diluted in PBS for 30 min and then washed three times with PBST (PBS supplemented with 0.2% Triton X-100) with 5 min intervals. To label the nucleus, samples were stained with DAPI (1:5000; D9542, Sigma Aldrich) in PBST supplemented with 1% BSA (A7906, Sigma Aldrich) for 1 hr at room temperature. Stained cells samples were washed three times with PBST and preserved in Vectashield (H-1000, Vector Laboratories, Burlingame, CA, USA). Fluorescence images were acquired using an Olympus FV1200 confocal microscope with 40X oil objective lens and 2X zoom factor. NIH ImageJ software was used for further adjustment and assembly of the acquired images.

#### Database searching and analysis of mass spectrometry data

MS/MS spectra were queried using the Comet search engine (Eng, Jahan, & Hoopmann, 2013) to search for corresponding proteins in Flybase (Gramates et al., 2017) and Uniprot (The UniProt, 2017). Common contaminant protein sequences from the Common Repository of Adventitious Proteins (cRAP) Database (ftp://ftp.thegpm.org/fasta/cRAP) were used to filter contaminating sequences. Searching was done with following parameters: tryptic digest, internal decoy peptides, the number of missed cleavages=2, precursor tolerance allowing for isotope offsets=20 ppm, a 1.00 fragment bin tolerance, static modification of 57.02 on cysteine, and variable modification of 16.00 on methionine. The acetylation, phosphorylation, and ubiquitination searches add variable modifications of 42.01 on lysine, 79.97 on serine/threonine/tyrosine, and 114.04 on lysine, respectively. The search results were then processed through the Trans-Proteomic Pipeline suite of tools version 4.8.0 (Keller, Eng, Zhang,

Li, & Aebersold, 2005) where the PeptideProphet tool (Keller, Nesvizhskii, Kolker, & Aebersold, 2002) was applied to calculate the probability that each search result is correct and the ProteinProphet tool (Nesvizhskii, Keller, Kolker, & Aebersold, 2003) was applied to infer protein identifications and their probabilities.

**Functional annotations and multiple sequence alignment of α-arrestin sequences**

The sequences of twelve *Drosophila* and six human α-arrestins were retrieved from the Uniprot database (UniProt Consortium, 2018). Domains and motifs including the PPxY motif were annotated based on sequences from Pfam version 31.0 (El-Gebali et al., 2019) and the eukaryotic linear motif (ELM) database (Dinkel et al., 2015). The sequences were subjected to the multiple-sequence alignment tool T-COFFEE (Notredame, Higgins, & Heringa, 2000) using default parameters. The output of T-COFFEE was applied to RAxML (version 8.2.11) (Stamatakis, 2014) to generate a consensus phylogenetic tree with 1,000 rapid bootstrapping using "-m PROTGAMMAWAGF" as the parameter.

**Checking the reproducibility of spectral counts among replicates**

If multiple proteins isoforms were detected, they were collapsed into a single gene. To avoid the divide-by-zero error, spectral counts of "0" were converted to a minimum non-zero value, "0.01". To examine the integrity and quality of spectral counts from the AP/MS, the average correlation coefficients (Pearson) of spectral counts from α-arrestins were calculated and plotted. At each cutoff of spectra counts from 1 to 15, only the PPIs with spectral counts that were the same or higher than the cutoff for all replicates were kept and used to calculate correlation coefficients between replicates. The resulting coefficients from the α-arrestin interactomes were then averaged and plotted. At the cutoff of 6 spectral counts, saturation of average correlation coefficients was observed and chosen as an optimal cutoff to filter the PPIs. Principal component analysis (PCA) of the filtered PPIs was conducted based on spectral counts (with a pseudo count 1 added) transformed into a $\log_2$ using the factoextra R package (version 1.0.7).

**Hierarchical clustering of high-confidence PPIs**

Hierarchical clustering based on $\log_2$ spectral counts (pseudo count 1 added) of high-confidence PPIs was conducted using the Pearson correlation as the clustering distance and Ward's method as the clustering method. Heatmaps were visualized through the ComplexHeatmap R package (version 2.6.2) (Gu, Eils, & Schlesner, 2016). Six clusters were identified for each species based on the results of hierarchical clustering; the PANTHER protein class overrepresentation test was performed for the proteins in each cluster (Thomas et al., 2003). False discovery rates (FDRs, Fisher's exact test) of indicated protein classes were ≤ 0.05 for all classes except for "GTPase-activating protein" in human (FDR < 0.133) and "GEFs" in *Drosophila* (FDR < 0.109), respectively. Interacting prey proteins from the positive PPIs were selectively labeled.

**Domain and motif analysis of bait and prey proteins**

For human and *Drosophila*, respectively, 53 and 65 short linear motifs in α-arrestins were annotated using the ELM database (Dinkel et al., 2015), and 423 and 546 protein domains in prey proteins were annotated using the Uniprot database (UniProt Consortium, 2018) (Table S4). To test for enrichment of protein domains, we implemented the Expression Analysis Systematic Explorer (EASE) score (Hosack, Dennis, Sherman, Lane, & Lempicki, 2003), which is calculated by subtracting one gene within the query domain and conducting a one-sided Fisher's exact test. Protein domains enriched in the interactomes of each α-arrestin (Benjamini-Hochberg FDR ≤ 0.05) were plotted using the ComplexHeatmap R package (version 2.6.2). Next, to see how reliable our filtered PPIs were, we utilized information about known affinities between domains and short linear motifs from the ELM database (Dinkel et al., 2015). Because the arrestin_N (Pfam ID : PF00339) and arrestin_C (Pfam Id : PF02752) domains in α-arrestins do not have known interactions with any of the short linear motifs in the ELM database (Dinkel et al., 2015), only the interactions between the short linear motifs in α-arrestins and protein domains in the interactome (prey proteins) were considered in this analysis. We found that 59 out of the 390 human PPIs and 64 out of the 740 *Drosophila* PPIs were supported by such known affinities (Table S4). One-sided Fisher's exact test was used to test the significance of the enrichment of the supported PPIs in the filtered PPI sets versus those in the unfiltered PPI sets (Figure 1D).

**Subcellular localizations of bait and prey proteins**

To search for annotated subcellular localizations of the proteins in the α-arrestin interactomes, we first obtained annotation files of cellular components (Gene Ontology (GO) : CC) for human and *Drosophila* from the Gene Ontology Consortium (Ashburner et al., 2000). From the annotations, we only utilized GO terms for 11 subcellular localizations (name of subcellular localization – GO term ID: Cytosol – GO:0005829; Plasma membrane – GO:0005886; Nucleus – GO:0005634; Mitochondrion – GO:0005739; Endoplasmic reticulum – GO:0005783; Golgi apparatus – GO:0005794; Cytoskeleton – GO:0005856; Peroxisome – GO:0005777; Lysosome – GO:0005764; Endosome – GO:0005768; Extracellular space – GO:0005615). If a protein was annotated to be localized in multiple locations, a weighted value (1/the number of multiple localizations) was assigned to each location. Finally, the relative frequencies of the subcellular localizations associated with the interacting proteins in the filtered PPIs were plotted for each α-arrestin (Figure S3B).

**Immunoblotting and co-immunoprecipitation Assays**

Cells were lysed in radioimmunoprecipitation assay (RIPA) buffer supplemented with protease inhibitor. For immunoblotting, the cell lysates were separated by 4-20% SDS-polyacrylamide gel electrophoresis (PAGE) and transferred to nitrocellulose membranes. After blocking membranes with 5% skim milk in Tris buffered Saline containing 0.1% Tween-20 (TBS-

T) for 1~2 hours (hr) at room temperature, the nitrocellulose membranes were incubated with appropriate primary antibodies overnight at 4°C and subsequently reacted with horseradish peroxidase (HRP)-conjugated secondary antibodies for 1 hr at room temperature. Bands were visualized using an enhanced chemiluminescence (ECL) detection system, West-Q Pico ECL Solution (W3652-02, GenDEPOT, Katy, TX, USA) or Fusion FX Spectra (Vilber, Marne-la-Vallée, France). For quantification of immunoblot results, the densities of target protein bands were analyzed with Image J.

For immunoprecipitation, the cell lysates (2 mg) were incubated with appropriate antibodies (1 µg) overnight at 4°C and precipitated with TrueBlot Anti-Rabbit Ig IP agarose beads (Rockland, Philadelphia, PA) for 2 hr at 4°C. The immunocomplexes were washed with chilled PBS three times and heated with 3x sample loading buffer containing ß-mercaptoethanol. The samples were separated by 6-8 % SDS-polyacrylamide gel electrophoresis (PAGE) and immunoblot was performed as described above.

The following antibodies were used for immunoblotting and co-immunoprecipitation assays: anti-TXNIP (#14715), anti-HDAC2 (#57156) ,anti-alpha Tubulin (#3873), anti-phospho-HDAC2 (#69238), anti-MTA1 (#5647), anti-MBD3 (#14540), anti-ATP6V1B2 (14617S), HRP-linked anti-rabbit IgG (7074S), and HRP-linked anti-mouse IgG (7076S) were obtained from Cell Signaling Technology (Beverly, MA); anti-H3ac (39139) was obtained from Active Motif (Carlsbad, CA); anti-ß-actin (GTX629630) was obtained from GeneTex; normal anti-rabbit IgG (sc-2027) and anti-GAPDH (sc-365062) were obtained from Santa Cruz Biotechnology (Dallas, TX); TrueBlot anti-rabbit IgG HRP (18-8816-31) was obtained from Rockland (Philadelphia, PA).

**Quantitative Reverse-transcription polymerase chain reaction (PCR)**

Total RNA was isolated using TRIzol reagent (#15596018, Invitrogen, Carlsbad, CA, USA; Thermo Fisher Scientific) and subjected to reverse transcription PCR (RT-PCR) with ReverTra Ace qPCR RT kit (#FSQ-101, Toyobo, Osaka, Japan) or GoScript RT-PCR system

(#A5001, Promega, Madison, WI, USA) according to the manufacturer's instructions. The mRNA expression levels of target genes were quantified using the CFX Opus 96 (Biorad, Hercules, CA) or Applied Biosystems QuantStudio 1 (Applied Biosystems, Foster city, CA) real-time PCR. AccuPower 2X GreenStar™ qPCR Master Mix (#K6251, Bioneer, Daejeon, Republic of Korea) or SYBR Green Realtime PCR Master Mix (#QPK-201, Toyobo, Osaka, Japan) were applied according to the manufacturer's protocols. The data normalized by GAPDH or alpha-tubulin mRNA levels and calculated using the ΔΔCt method (Hellemans, Mortier, De Paepe, Speleman, & Vandesompele, 2007). The primers used for qRT-PCR analysis are summarized in Table S12.

**PCA of ATAC- and RNA-seq data**

For ATAC-seq, normalized read counts derived from the diffBind R package (version 3.0.15) (Ross-Innes et al., 2012) were transformed into a $\log_2$ function. Batch effect corrections were done using the limma R package (version 3.46.0) (Ritchie et al., 2015). For RNA-seq, counts per million mapped reads (CPM) were also processed in the same manner. For PCA, 2,000 features with the highest variance across samples were extracted and utilized. Plots of principal components 1 and 2 were generated by the factoextra R package (version 1.0.7).

**Functional signatures of repressed genes upon TXNIP depletion**

Genes that exhibited decreased chromatin accessibility at their promoter and decreased RNA expression upon TXNIP knockdown (Table S11) were selected based on the following criteria: 1. $\log_2$ (RNA level in siTXNIP-treated cells/RNA level in siCon-treated cells) (hereafter, siTXNIP/siCon) $\leq$ -1; 2. $\log_2$ (siTXNIP/siCon) of ACRs in the promoter region $\leq$ -1 (If there are multiple ACRs in the promoter region, the one with the highest ATAC-seq signal was selected) or $\log_2$ mean (siTXNIP/siCon) of all ACRs in the promoter region $\leq$ -1. Enrichment analysis of the GO terms in the gene set was performed by g:Profiler (Raudvere et al., 2019).

Top 10 enriched terms from the biological process and molecular functions categories were plotted (Figure 4G).

**Immunofluorescence of HDAC2 and TXNIP**

HeLa cells were cultured in 6-well plates with cover slips in each well (1.5 $x10^4$ cells/well). After cells were incubated overnight in Opti-MEM, TXNIP knockdown was induced by transfection of siRNA at a concentration of 100 nM. Following 48 hr of transfection, the cells were washed twice with PBS and then fixed with 100% ice-cold methanol for 10 min at -20˚C. After rinsing three with PBSTw (PBS containing 0.1% Tween 20), the cells were blocked with 3% BSA in PBS and incubated for 45 min at room temperature. Next, cells were incubated with the primary antibody for 150 min followed by the secondary antibody for 60 min in the dark. For co-staining with a second primary antibody, the blocking step followed by the primary and secondary antibody incubation steps were repeated. All of the antibodies were diluted in antibody dilution buffer (1% BSA in PBS). Information of the antibodies are listed in "antibody" section in STAR Method. The cover slips were rinsed three times with PBSTw and then mounted with VECTASHIELD Antifade Mounting Medium containing DAPI (Vector Laboratories, Newark, CA, USA) according to the manufacturer's instructions. The fluorescence was visualized with a Nikon C2 Si-plus confocal microscope. Fluorescence images were observed under a ZEISS confocal microscope (LSM5; Carl Zeiss, Jena, Germany), and the integrated densities of fluorescence were analyzed using ImageJ program.

### Supplementary figures

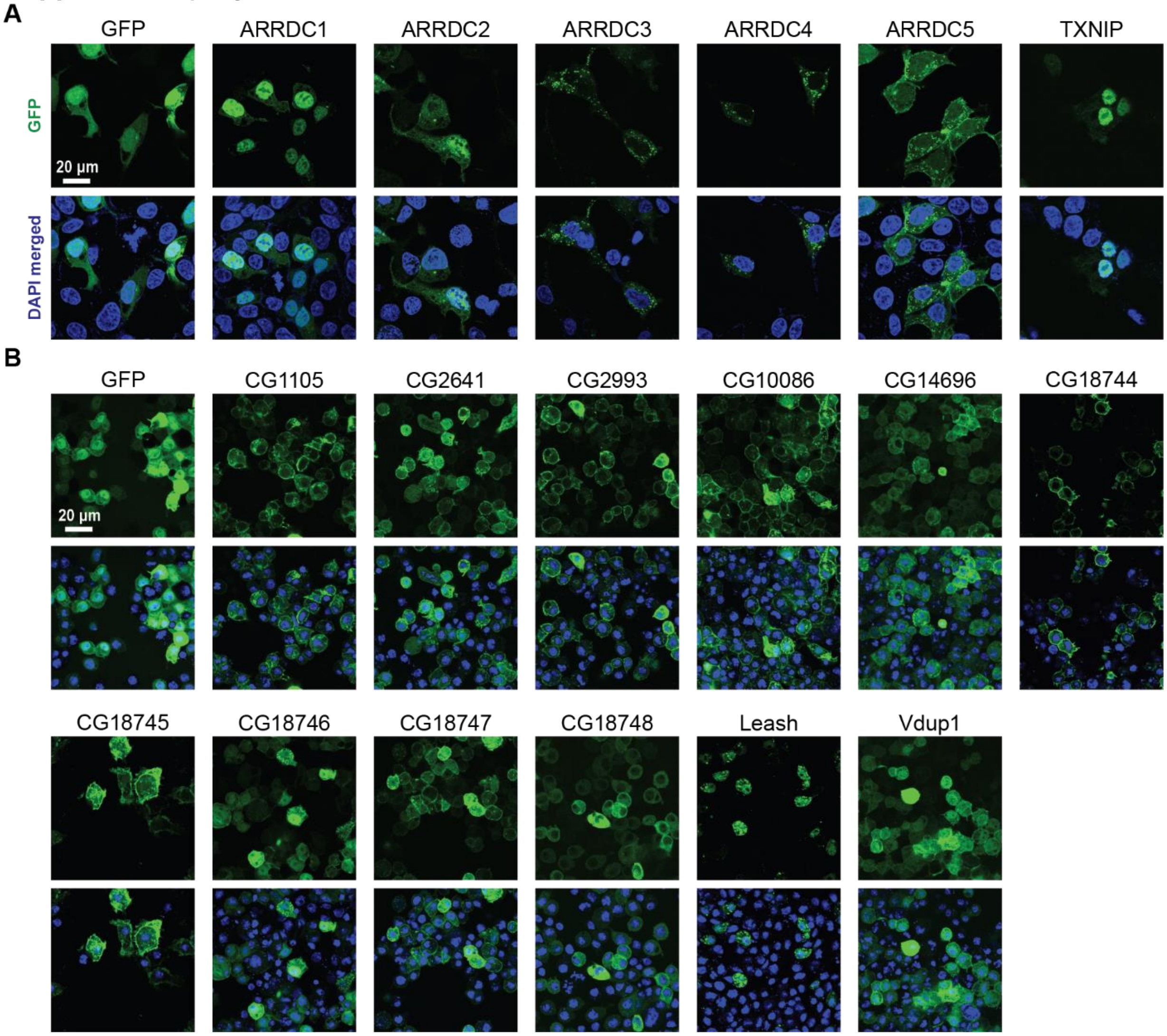


**Figure S1. Fluorescence images showing HEK293 and S2R+ cells stably expressing GFP-tagged α-arrestins**

Representative images of HEK293 **(A)** and S2R+ cells **(B)** stably expressing GFP-tagged α-arrestins.

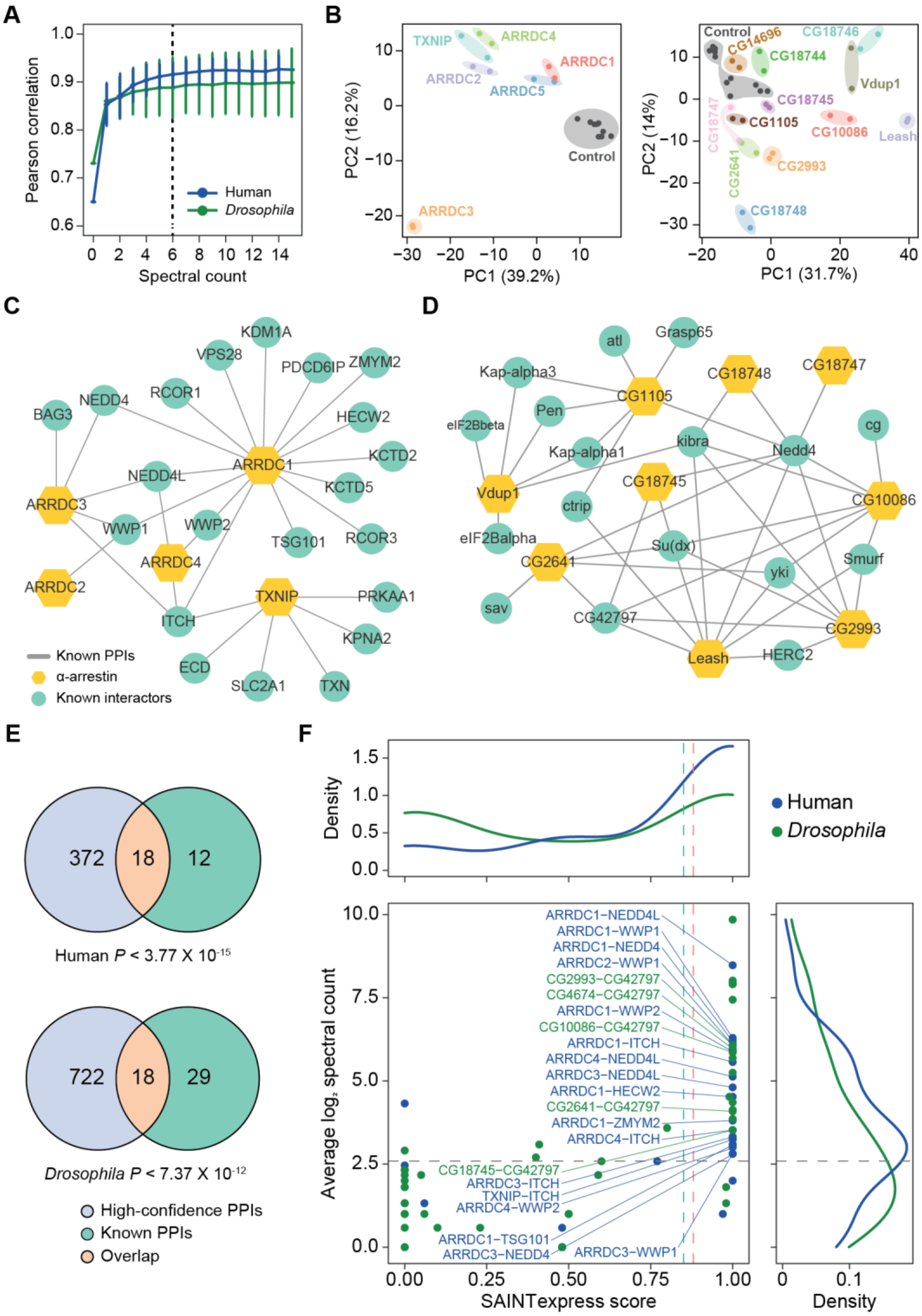

A
Pearson correlation
Spectral count
Human
Drosophila
B
TXNIP
ARRDC4
ARRDC1
ARRDC2
ARRDC5
Control
ARRDC3
PC2 (16.2%)
PC1 (39.2%)
CG14696
CG18746
CG18744
Vdup1
CG18745
CG18747
CG1105
CG10086
Leash
CG2641
CG2993
CG18748
PC2 (14%)
PC1 (31.7%)
C
Known PPIs
α-arrestin
Known interactors
D
E
372
18
12
Human P < 3.77 X 10-15
722
18
29
Drosophila P < 7.37 X 10-12
High-confidence PPIs
Known PPIs
Overlap
F
Density
Human
Drosophila
Average log2 spectral count
SAINTexpress score
ARRDC1−NEDD4L
ARRDC1−WWP1
ARRDC1−NEDD4
ARRDC2−WWP1
CG2993−CG42797
CG4674−CG42797
ARRDC1−WWP2
CG10086−CG42797
ARRDC1−ITCH
ARRDC4−NEDD4L
ARRDC3−NEDD4L
ARRDC1−HECW2
CG2641−CG42797
ARRDC1−ZMYM2
ARRDC4−ITCH
CG18745−CG42797
ARRDC3−ITCH
TXNIP−ITCH
ARRDC4−WWP2
ARRDC1−TSG101
ARRDC3−NEDD4
ARRDC3−WWP1

**Figure S2. AP/MS data offer high reproducibility and expand the PPIs associated with α-arrestins while also reaffirming known interactions**

**(A)** Average Pearson correlation coefficients of $\log_2$ spectral counts between replicates of AP/MS of each α-arrestin at varying cutoffs are shown (mean ± standard deviation(sd)). The cutoff used in this study, 6, is shown as a dashed line. **(B)** PCA plots based on $\log_2$ spectral counts of high-confidence PPIs for human (left) and *Drosophila* (right) are shown. **(C and D)** PPI network of α-arrestins that were previously reported and utilized as the positive PPI sets (Table S2A and S2C) for human **(C)** and *Drosophila* **(D)**. **(E)** Venn diagram showing the intersection of the high-confidence PPIs and positive PPI sets associated with α-arrestins for human (top, Table S2A) and *Drosophila* (bottom, Table S2C). *P* values were determined via the hypergeometric test. **(F)** Distribution of SAINTexpress scores and average spectral counts ($\log_2$) of the positive PPIs (Table S2A and C) are shown and density plots for each axis are also plotted. The positive PPIs that are also included in the high-confidence PPIs are selectively labeled.

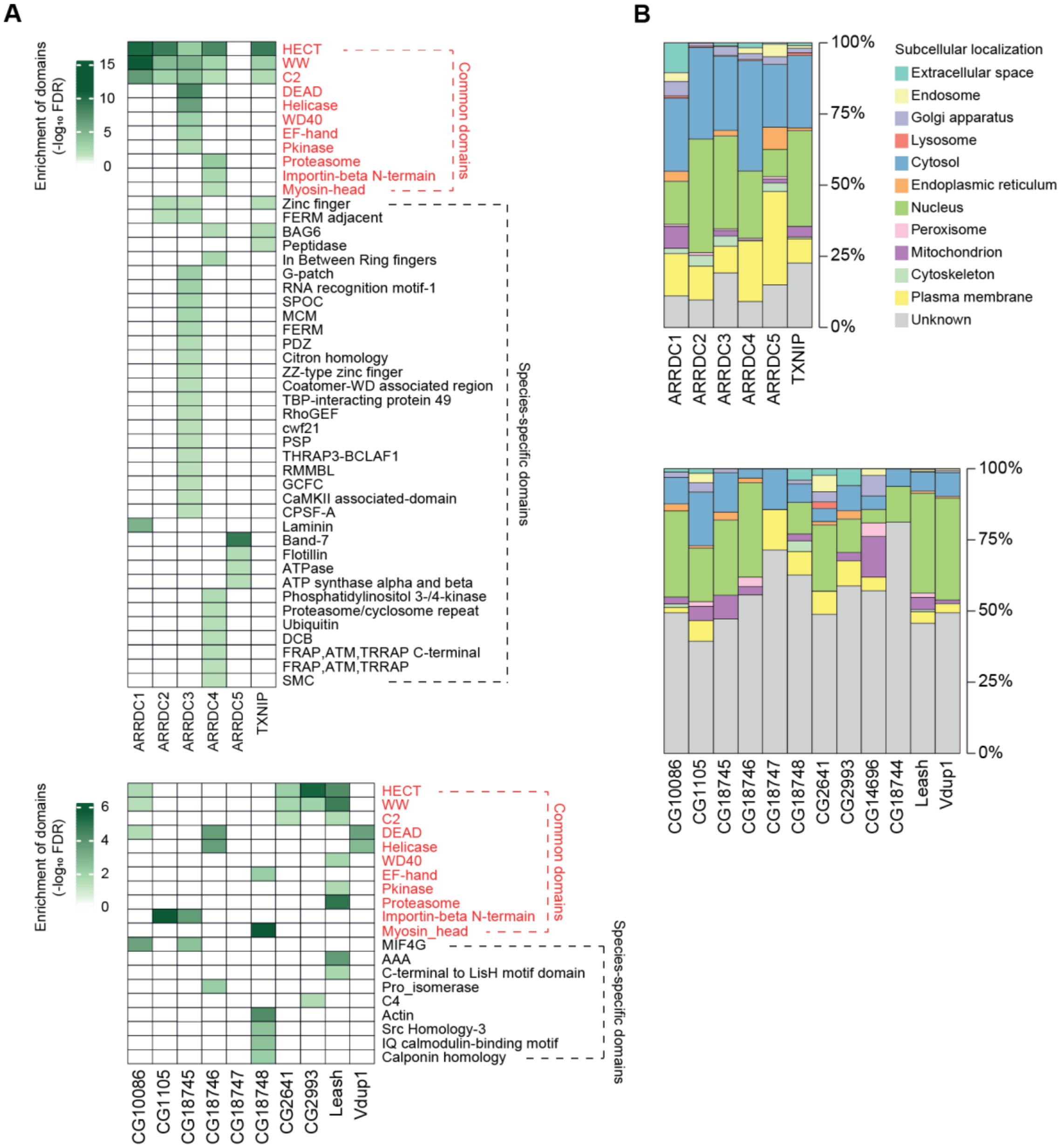
A
Enrichment of domains (-log10 FDR)
15
10
5
0
HECT
WW
C2
DEAD
Helicase
WD40
EF-hand
Pkinase
Proteasome
Importin-beta N-termain
Myosin-head
Common domains
Zinc finger
FERM adjacent
BAG6
Peptidase
In Between Ring fingers
G-patch
RNA recognition motif-1
SPOC
MCM
FERM
PDZ
Citron homology
ZZ-type zinc finger
Coatomer-WD associated region
TBP-interacting protein 49
RhoGEF
cwf21
PSP
THRAP3-BCLAF1
RMMBL
GCFC
CaMKII associated-domain
CPSF-A
Laminin
Band-7
Flotillin
ATPase
ATP synthase alpha and beta
Phosphatidylinositol 3-/4-kinase
Proteasome/cyclosome repeat
Ubiquitin
DCB
FRAP,ATM,TRRAP C-terminal
FRAP,ATM,TRRAP
SMC
Species-specific domains
ARRDC1
ARRDC2
ARRDC3
ARRDC4
ARRDC5
TXNIP
Enrichment of domains (-log10 FDR)
6
4
2
0
HECT
WW
C2
DEAD
Helicase
WD40
EF-hand
Pkinase
Proteasome
Importin-beta N-termain
Myosin_head
Common domains
MIF4G
AAA
C-terminal to LisH motif domain
Pro_isomerase
C4
Actin
Src Homology-3
IQ calmodulin-binding motif
Calponin homology
Species-specific domains
CG10086
CG1105
CG18745
CG18746
CG18747
CG18748
CG2641
CG2993
Leash
Vdup1
B
100%
75%
50%
25%
0%
ARRDC1
ARRDC2
ARRDC3
ARRDC4
ARRDC5
TXNIP
Subcellular localization
Extracellular space
Endosome
Golgi apparatus
Lysosome
Cytosol
Endoplasmic reticulum
Nucleus
Peroxisome
Mitochondrion
Cytoskeleton
Plasma membrane
Unknown
100%
75%
50%
25%
0%
CG10086
CG1105
CG18745
CG18746
CG18747
CG18748
CG2641
CG2993
CG14696
CG18744
Leash
Vdup1

**Figure S3. Protein domains and subcellular localization of α-arrestin interactomes**

**(A)** Protein domains enriched in the interactome of each α-arrestin for human (top) and *Drosophila* (bottom) are shown. The significance of the enrichment test ($-\log_{10}$ FDR) is indicated in shades of green, as depicted in the legend. SPOC, spen paralogue and orthologue C-terminal; MCM, minichromosome maintenance protein complex; FDRM, F for 4.1 protein, E for ezrin, R for radixin and M for moesin; TBP, TATA binding protein; GEF, guanine nucleotide exchange factor; THRAP3, thyroid hormone receptor-associated protein 3; BCLAF1, Bcl-2-associated transcription factor1; RMMBL, RNA metabolizing metallo beta lactamase; CaMKII, C-terminus of the Calcium/calmodulin dependent protein kinases II; CPSF, cleavage and polyadenylation specificity factor; DCB, dimerization and cyclophilin-binding domain; FRAP, FKBP12-rapamycin complex-associated protein; ATM, ataxia telangiectasia mutant; THRAP, transformation/transcription domain associated proteins; MIF4G, middle domain of eukaryotic initiation factor 4G; AAA, ATPase family associated with various cellular activities; C4, C-terminal tandem repeated domain in type 4 procollagen; SMC, structural maintenance of chromosomes. **(B)** Subcellular localizations of the interactome of each α-arrestin for human (top) and *Drosophila* (bottom).

A

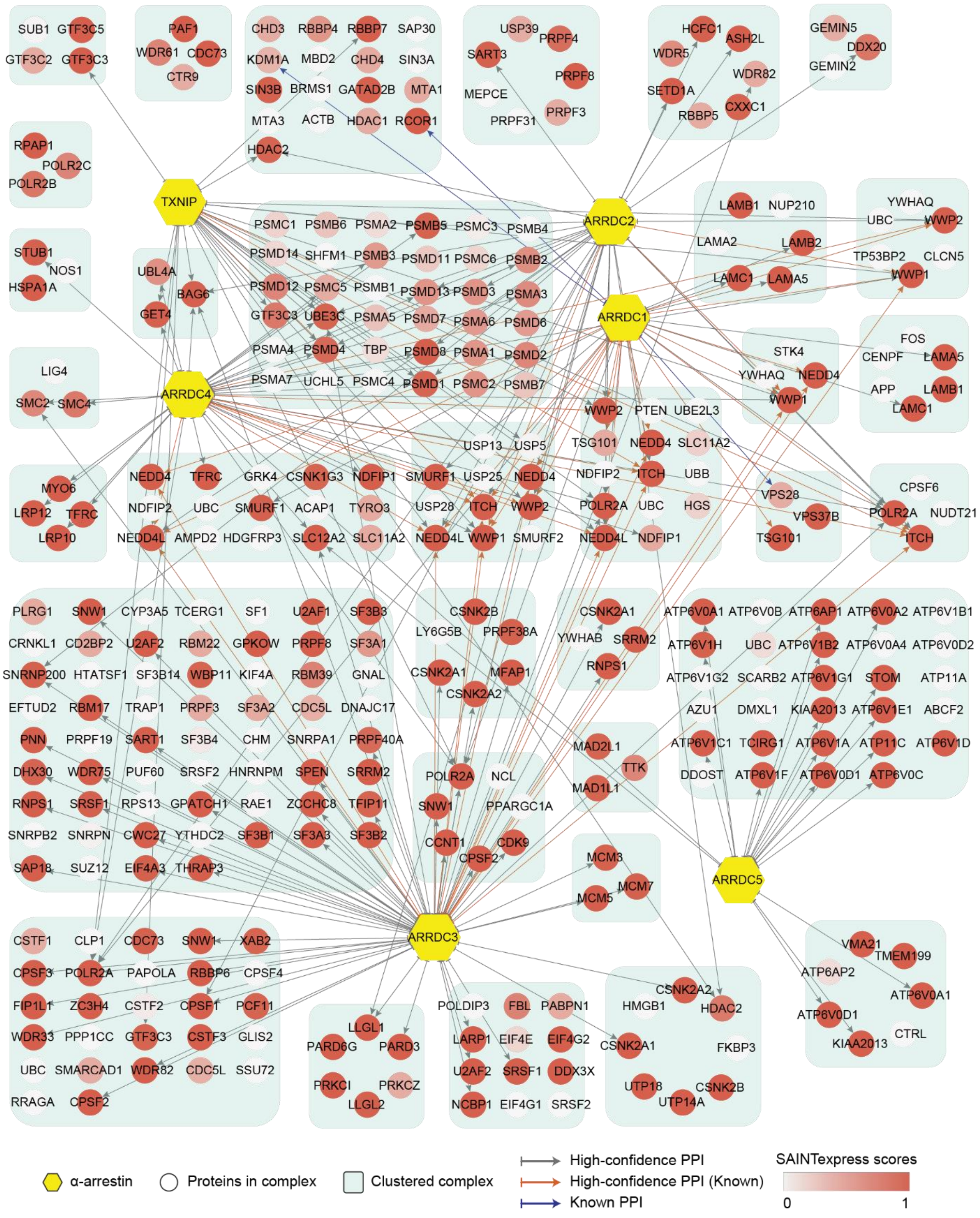

SUB1 GTF3C5 GTF3C2 GTF3C3
PAF1 WDR61 CDC73 CTR9
CHD3 RBBP4 RBBP7 SAP30 KDM1A MBD2 CHD4 SIN3A SIN3B BRMS1 GATAD2B MTA1 MTA3 ACTB HDAC1 RCOR1 HDAC2
USP39 PRPF4 SART3 PRPF8 MEPCE PRPF31 PRPF3
HCFC1 ASH2L WDR5 WDR82 SETD1A RBBP5 CXXC1
GEMIN5 DDX20 GEMIN2
RPAP1 POLR2C POLR2B
TXNIP
ARRDC2
PSMC1 PSMB6 PSMA2 PSMB5 PSMC3 PSMB4
PSMD14 SHFM1 PSMB3 PSMD11 PSMC6 PSMB2
PSMD12 PSMC5 PSMB1 PSMD13 PSMD3 PSMA3
GTF3C3 UBE3C PSMA5 PSMD7 PSMA6 PSMD6
PSMA4 PSMD4 TBP PSMD8 PSMA1 PSMD2
PSMA7 UCHL5 PSMC4 PSMD1 PSMC2 PSMB7
LAMB1 NUP210 LAMA2 LAMB2 LAMC1 LAMA5
YWHAQ UBC WWP2 TP53BP2 CLCN5 WWP1
STUB1 NOS1 HSPA1A
UBL4A BAG6 GET4
ARRDC1
LIG4 SMC2 SMC4
ARRDC4
STK4 YWHAQ NEDD4 WWP1
FOS CENPF LAMA5 APP LAMB1 LAMC1
WWP2 PTEN UBE2L3 TSG101 NEDD4 SLC11A2 NDFIP2 ITCH UBB POLR2A UBC HGS NEDD4L NDFIP1
USP13 USP5 SMURF1 USP25 NEDD4 USP28 ITCH WWP2 NEDD4L WWP1 SMURF2
MYO6 LRP12 TFRC LRP10
NEDD4 TFRC GRK4 CSNK1G3 NDFIP1 NDFIP2 UBC SMURF1 ACAP1 TYRO3 NEDD4L AMPD2 HDGFRP3 SLC12A2 SLC11A2
VPS28 VPS37B TSG101
CPSF6 POLR2A NUDT21 ITCH
PLRG1 SNW1 CYP3A5 TCERG1 SF1 U2AF1 SF3B3
CRNKL1 CD2BP2 U2AF2 RBM22 GPKOW PRPF8 SF3A1
SNRNP200 HTATSF1 SF3B14 WBP11 KIF4A RBM39 GNAL
EFTUD2 RBM17 TRAP1 PRPF3 SF3A2 CDC5L DNAJC17
PNN PRPF19 SART1 SF3B4 CHM SNRPA1 PRPF40A
DHX30 WDR75 PUF60 SRSF2 HNRNPM SPEN SRRM2
RNPS1 SRSF1 RPS13 GPATCH1 RAE1 ZCCHC8 TFIP11
SNRPB2 SNRPN CWC27 YTHDC2 SF3B1 SF3A3 SF3B2
SAP18 SUZ12 EIF4A3 THRAP3
CSNK2B LY6G5B PRPF38A CSNK2A1 MFAP1 CSNK2A2
CSNK2A1 YWHAB SRRM2 RNPS1
ATP6V0A1 ATP6V0B ATP6AP1 ATP6V0A2 ATP6V1B1
ATP6V1H UBC ATP6V1B2 ATP6V0A4 ATP6V0D2
ATP6V1G2 SCARB2 ATP6V1G1 STOM ATP11A
AZU1 DMXL1 KIAA2013 ATP6V1E1 ABCF2
ATP6V1C1 TCIRG1 ATP6V1A ATP11C ATP6V1D
DDOST ATP6V1F ATP6V0D1 ATP6V0C
POLR2A NCL SNW1 PPARGC1A CCNT1 CDK9 CPSF2
MAD2L1 TTK MAD1L1
MCM3 MCM7 MCM5
ARRDC5
ARRDC3
CSTF1 CLP1 CDC73 SNW1 XAB2
CPSF3 POLR2A PAPOLA RBBP6 CPSF4
FIP1L1 ZC3H4 CSTF2 CPSF1 PCF11
WDR33 PPP1CC GTF3C3 CSTF3 GLIS2
UBC SMARCAD1 WDR82 CDC5L SSU72
RRAGA CPSF2
LLGL1 PARD6G PARD3 PRKCI PRKCZ LLGL2
POLDIP3 FBL PABPN1 LARP1 EIF4E EIF4G2 U2AF2 SRSF1 DDX3X NCBP1 EIF4G1 SRSF2
CSNK2A2 HMGB1 HDAC2 CSNK2A1 FKBP3 UTP18 CSNK2B UTP14A
VMA21 TMEM199 ATP6AP2 ATP6V0A1 ATP6V0D1 CTRL KIAA2013
α-arrestin
Proteins in complex
Clustered complex
High-confidence PPI
High-confidence PPI (Known)
Known PPI
SAINTexpress scores
0
1

B

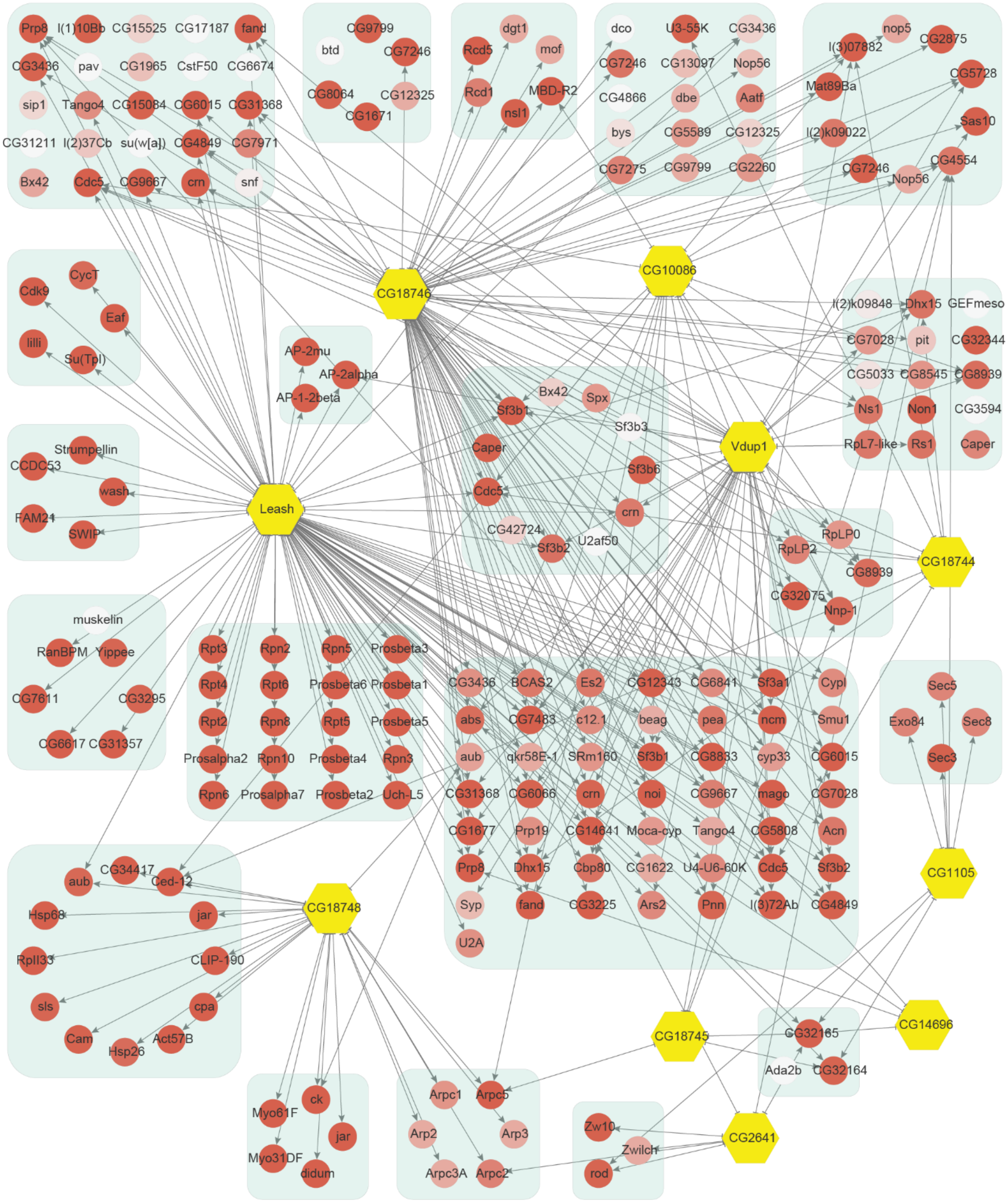

Prp8
l(1)10Bb
CG15525
CG17187
fand
CG3436
pav
CG1965
CstF50
CG6674
sip1
Tango4
CG15084
CG6015
CG31368
CG31211
l(2)37Cb
su(w[a])
CG4849
CG7971
Bx42
Cdc5
CG9667
crn
snf
CG9799
btd
CG7246
CG8064
CG12325
CG1671
dgt1
Rcd5
mof
Rcd1
MBD-R2
nsl1
dco
U3-55K
CG3436
CG7246
CG13097
Nop56
CG4866
dbe
Aatf
bys
CG5589
CG12325
CG7275
CG9799
CG2260
nop5
l(3)07882
CG2875
Mat89Ba
CG5728
l(2)k09022
Sas10
CG7246
CG4554
Nop56
CycT
Cdk9
Eaf
lilli
Su(Tpl)
CG18746
CG10086
AP-2mu
AP-2alpha
AP-1-2beta
l(2)k09848
Dhx15
GEFmeso
CG7028
pit
CG32344
CG5033
CG8545
CG8939
Ns1
Non1
CG3594
RpL7-like
Rs1
Caper
Bx42
Spx
Sf3b1
Sf3b3
Caper
Vdup1
Sf3b6
Strumpellin
CCDC53
wash
FAM21
SWIP
Leash
Cdc5
crn
CG42724
Sf3b2
U2af50
RpLP0
RpLP2
CG18744
CG8939
CG32075
Nnp-1
muskelin
RanBPM
Yippee
CG7611
CG3295
CG6617
CG31357
Rpt3
Rpn2
Rpn5
Prosbeta3
Rpt4
Rpt6
Prosbeta6
Prosbeta1
Rpt2
Rpn8
Rpt5
Prosbeta5
Prosalpha2
Rpn10
Prosbeta4
Rpn3
Rpn6
Prosalpha7
Prosbeta2
Uch-L5
CG3436
BCAS2
Es2
CG12343
CG6841
Sf3a1
Cypl
abs
CG7483
c12.1
beag
pea
ncm
Smu1
aub
qkr58E-1
SRm160
Sf3b1
CG8833
cyp33
CG6015
CG31368
CG6066
crn
noi
CG9667
mago
CG7028
CG1677
Prp19
CG14641
Moca-cyp
Tango4
CG5808
Acn
Prp8
Dhx15
Cbp80
CG1622
U4-U6-60K
Cdc5
Sf3b2
Syp
fand
CG3225
Ars2
Pnn
l(3)72Ab
CG4849
U2A
Sec5
Exo84
Sec8
Sec3
CG1105
aub
CG34417
Ced-12
Hsp68
jar
CG18748
RpII33
CLIP-190
sls
cpa
Cam
Act57B
Hsp26
CG18745
CG32165
CG14696
Ada2b
CG32164
ck
Myo61F
jar
Myo31DF
didum
Arpc1
Arpc5
Arp2
Arp3
Arpc3A
Arpc2
Zw10
Zwilch
rod
CG2641

**Figure S4. Network of high-confidence PPIs between α-arrestins and individual proteins of its associated protein complexes**

In addition to interaction between α-arrestins and protein complexes in Figure 2, PPIs between α-arrestins and individual components of its associated protein complexes are depicted for human **(A)** and *Drosophila* **(B)**. "High-confidence PPI (Known)" refers to previously reported interactions (Table S2A and C) that are also present in the high-confidence PPIs, while "Known PPI" pertains to those previously reported but absent from our collection.

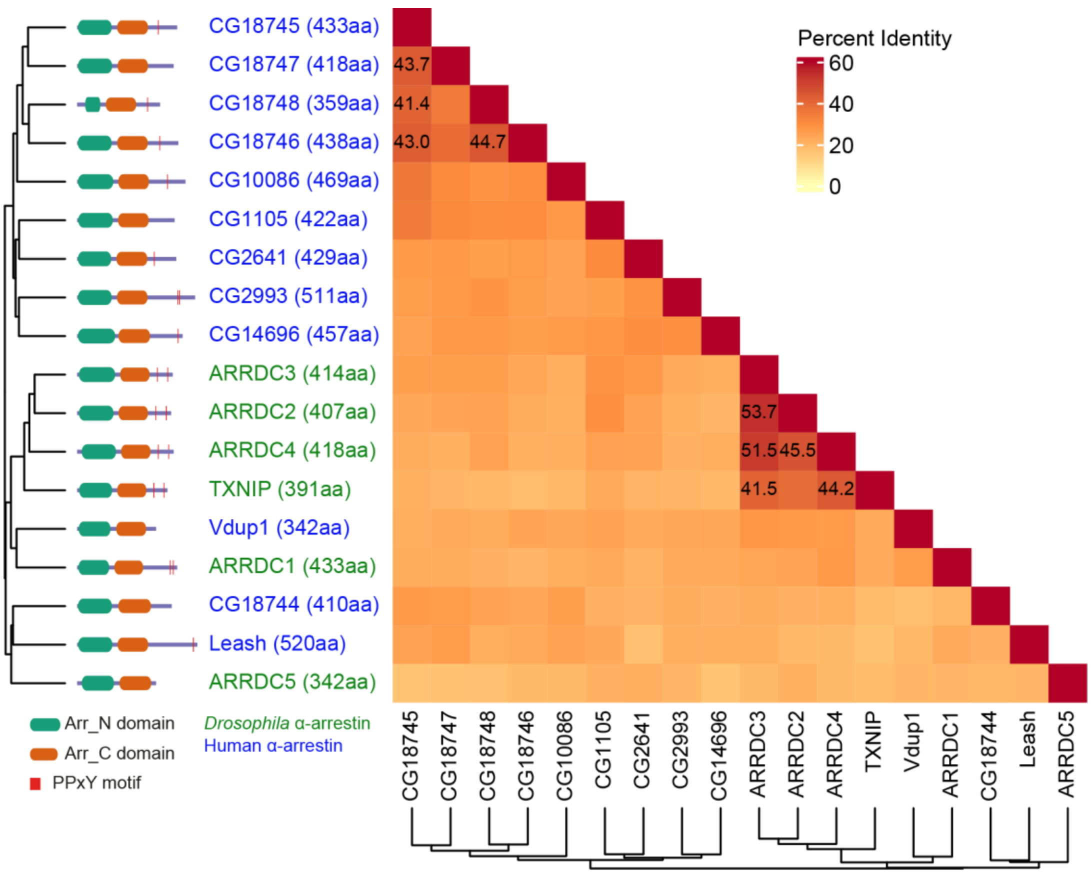


**Figure S5. Protein sequence homology of α-arrestins from human and *Drosophila***

Heatmap illustrates percent identity of protein sequences of α-arrestins from human and *Drosophila*.

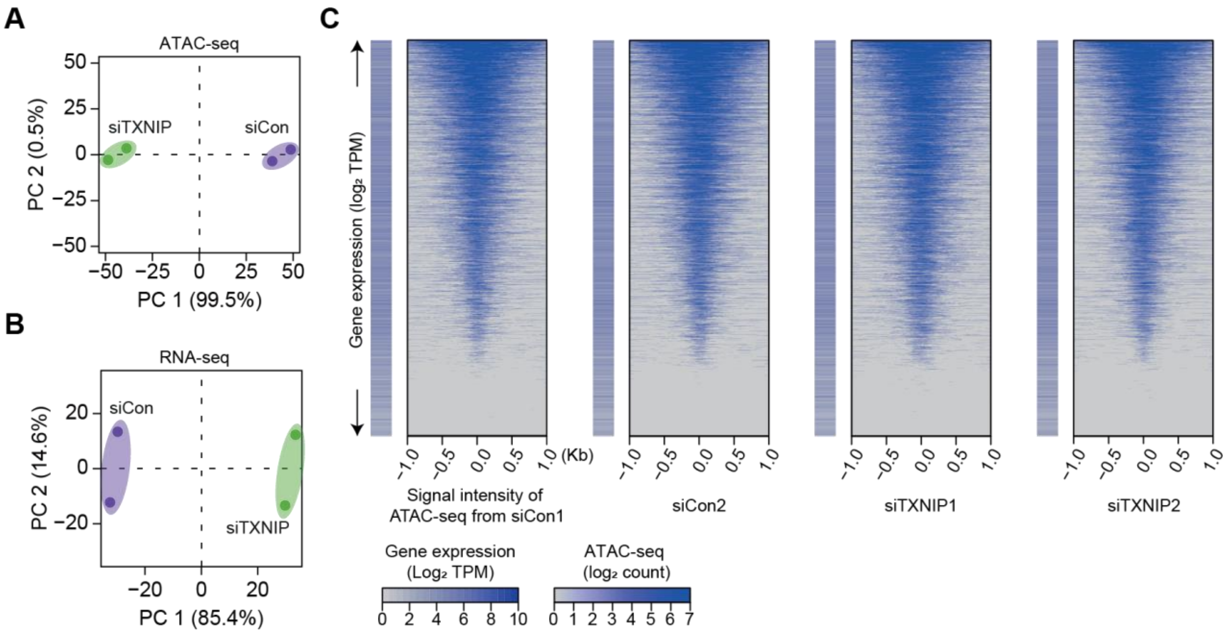


**Figure S6. High-throughput sequencing data are highly reproducible and ATAC-seq reads exhibit a typical pattern of strong enrichment around TSSs of expressed genes**

**(A and B)** PCA plots of ATAC- **(A)** and RNA-seq **(B)** results based on batch-corrected $\log_2$ counts and CPM, respectively. Numbers in parentheses are percentages of explained variance for the corresponding PCs. **(C)** Heatmaps of ATAC-seq read counts (read counts have been transformed into a $\log_2$ function and corrected for batch effects) in regions surrounding TSSs along with $\log_2$ transcript per million mapped reads (TPM, see “Materials and Methods”) for genes having the corresponding TSS are plotted for each sample.

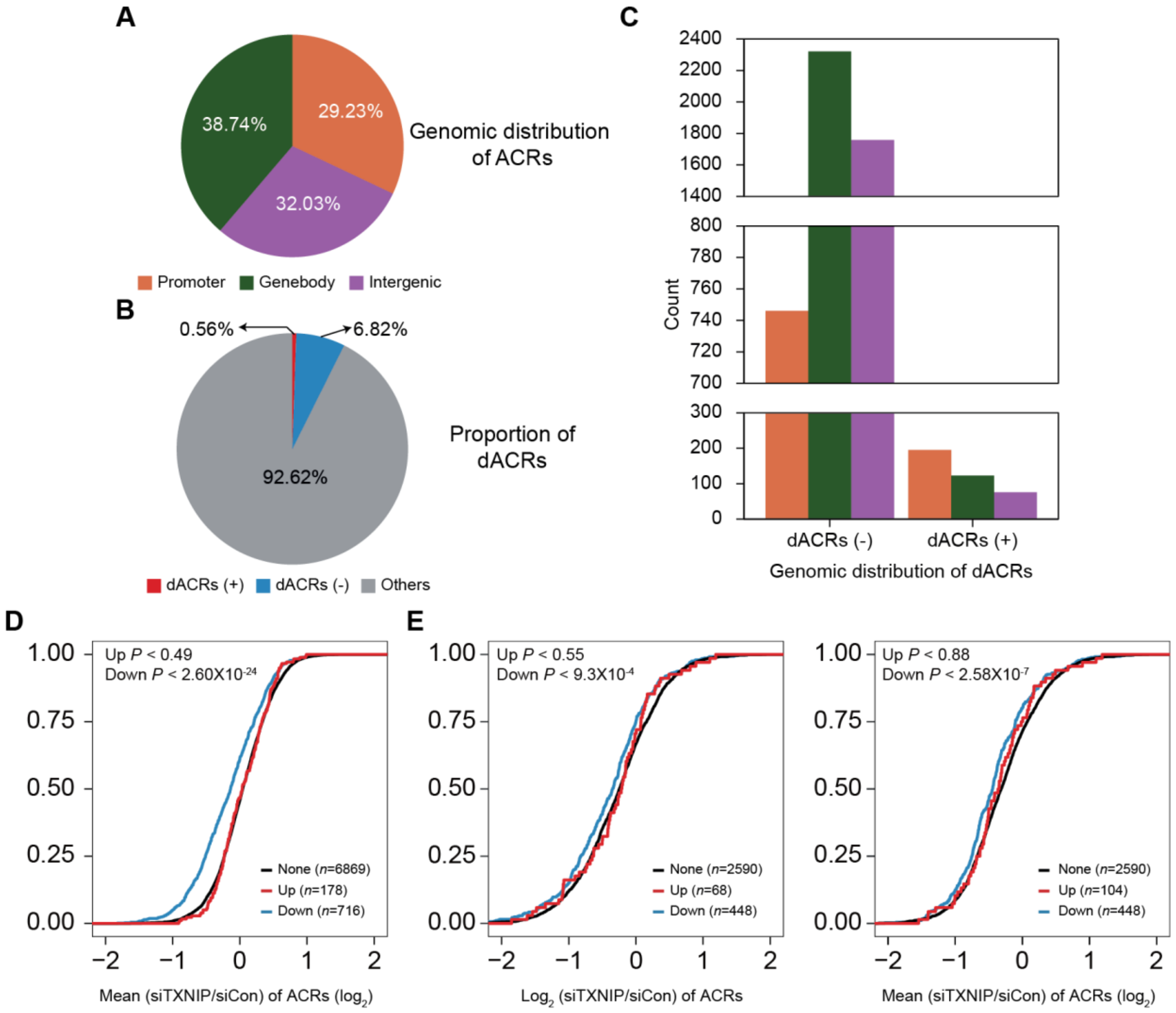
A
38.74%
29.23%
32.03%
Genomic distribution of ACRs
Promoter
Genebody
Intergenic
B
0.56%
6.82%
92.62%
Proportion of dACRs
dACRs (+)
dACRs (-)
Others
C
Count
dACRs (-)
dACRs (+)
Genomic distribution of dACRs
D
Up P < 0.49
Down P < 2.60X10-24
None (n=6869)
Up (n=178)
Down (n=716)
Mean (siTXNIP/siCon) of ACRs (log2)
E
Up P < 0.55
Down P < 9.3X10-4
None (n=2590)
Up (n=68)
Down (n=448)
Log2 (siTXNIP/siCon) of ACRs
Up P < 0.88
Down P < 2.58X10-7
None (n=2590)
Up (n=104)
Down (n=448)
Mean (siTXNIP/siCon) of ACRs (log2)

**Figure S7. Genomic locations of ACRs and association between chromatin accessibility and transcriptional activity**

**(A)** Genomic locations of 70,746 consensus ACRs identified from ATAC-seq analysis. **(B)** Composition of dACRs(-), dACRs(+), and other ACRs ("others", not significantly changed) under the TXNIP knockdown condition compared to the control. **(C)** Genomic locations of 4,825 dACRs(-) and 394 dACRs(+) are depicted. Colors in the bar plot have the same symbolism as in **(A)**. **(D)** Cumulative distribution function (CDF) of mean changes in accessibility of all ACRs located in gene promoters. The genes were categorized into three groups ("None", "Down", and "Up") as explained in Figure 4F. *P* values on the left upper corner were calculated with the one-sided Kolmogorov-Smirnov (KS) test, which compares "Up" or "Down" groups to the "None" group. **(E)** CDF of changes in accessibility of ACRs located in gene bodies. Changes in accessibility of ACRs whose intensity is highest among all ACRs located in gene bodies are depicted on the left and mean changes in accessibility of all ACRs located in gene bodies are depicted on the right. *P* values on the upper left corners are calculated in the same manner as in **(D)**.

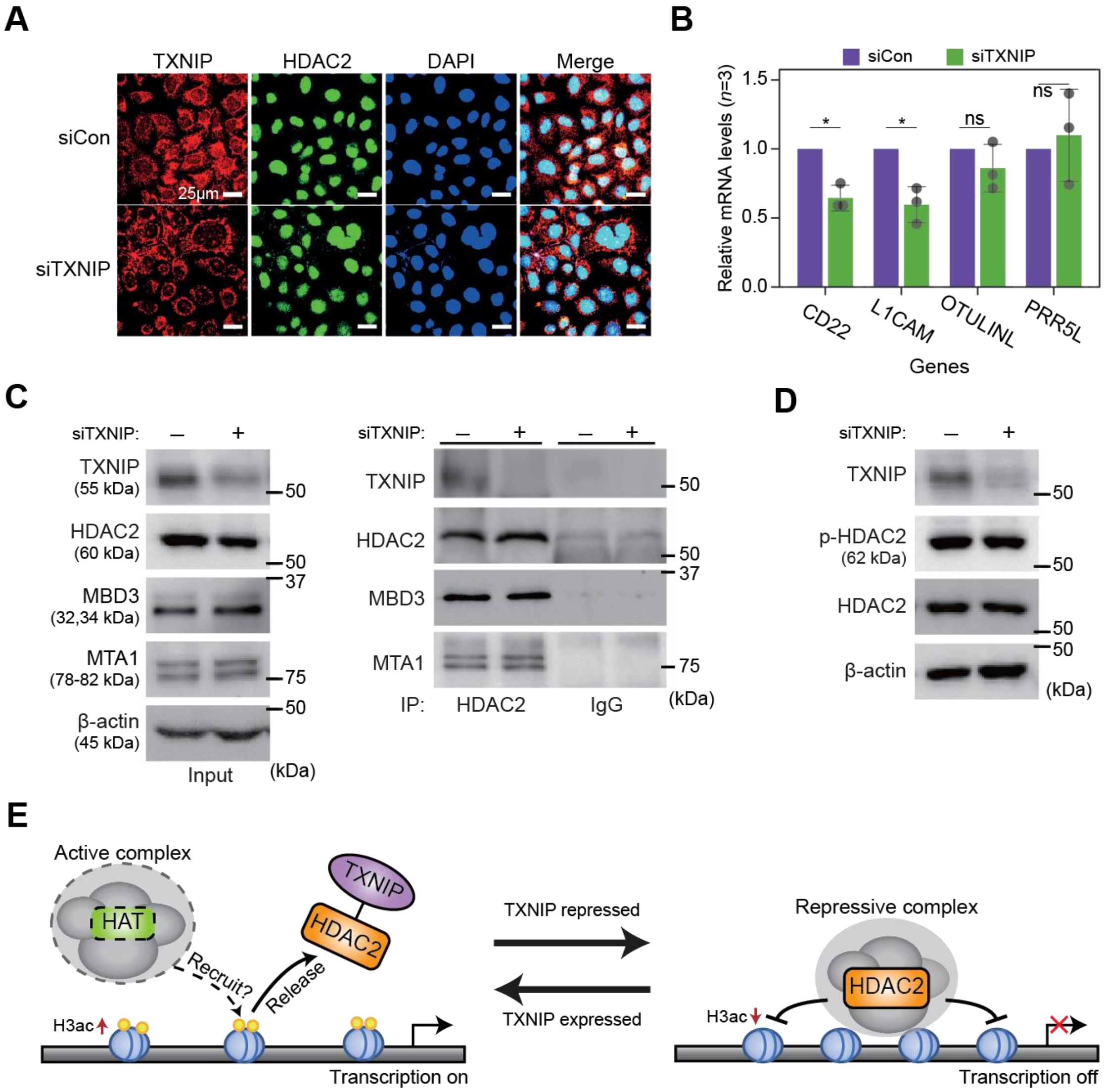

A
TXNIP
HDAC2
DAPI
Merge
siCon
siTXNIP
25μm
B
siCon
siTXNIP
Relative mRNA levels (n=3)
1.5
1.0
0.5
0.0
*
*
ns
ns
CD22
L1CAM
OTULINL
PRR5L
Genes
C
siTXNIP:
TXNIP (55 kDa)
HDAC2 (60 kDa)
MBD3 (32,34 kDa)
MTA1 (78-82 kDa)
β-actin (45 kDa)
Input
(kDa)
TXNIP
HDAC2
MBD3
MTA1
IP: HDAC2 IgG
50
37
75
D
siTXNIP:
TXNIP
p-HDAC2 (62 kDa)
HDAC2
β-actin
(kDa)
E
Active complex
HAT
Recruit?
Release
TXNIP
HDAC2
H3ac
Transcription on
TXNIP repressed
TXNIP expressed
Repressive complex
HDAC2
H3ac
Transcription off

**Figure S8. TXNIP might play role in transcriptional regulation independent of known factors**

**(A)** Representative immunofluorescence images of TXNIP and HDAC2 after HeLa cells were transfected with either siCon or siTXNIP for 48 hr (magnification ×600); TXNIP (red), HDAC2 (green), and DAPI (blue). **(B)** RT-qPCR results of four target genes whose RNA expression and chromatin accessibility in their promoters, quantified using high-throughput sequencing data, were observed to be strongly repressed in HeLa cell. Data are presented as the mean ± sd, n=3). Gray dots depict actual values of each experiment. * $P$ < 0.05, ns: not significant (two-sided paired Student's t-test). **(C)** Co-IP assay showing interaction of HDAC2 with TXNIP, MBD3, and MTA1 proteins. Lysates from HeLa cells were treated with either control siRNA (siCon) or TXNIP siRNA (siTXNIP) for 48 hours and subjected to immunoprecipitation and immunoblotting with the indicated antibodies. **(D)** Immunoblot analysis shows changes in the expression levels of phosphorylated HDAC2 (p-HDAC2) and HDAC2 protein in TXNIP knockdown HeLa cells. **(C-D)** β-actin was used as negative control. **(E)** A proposal model for the transcriptional regulation of target genes by interaction between TXNIP and HDAC2.

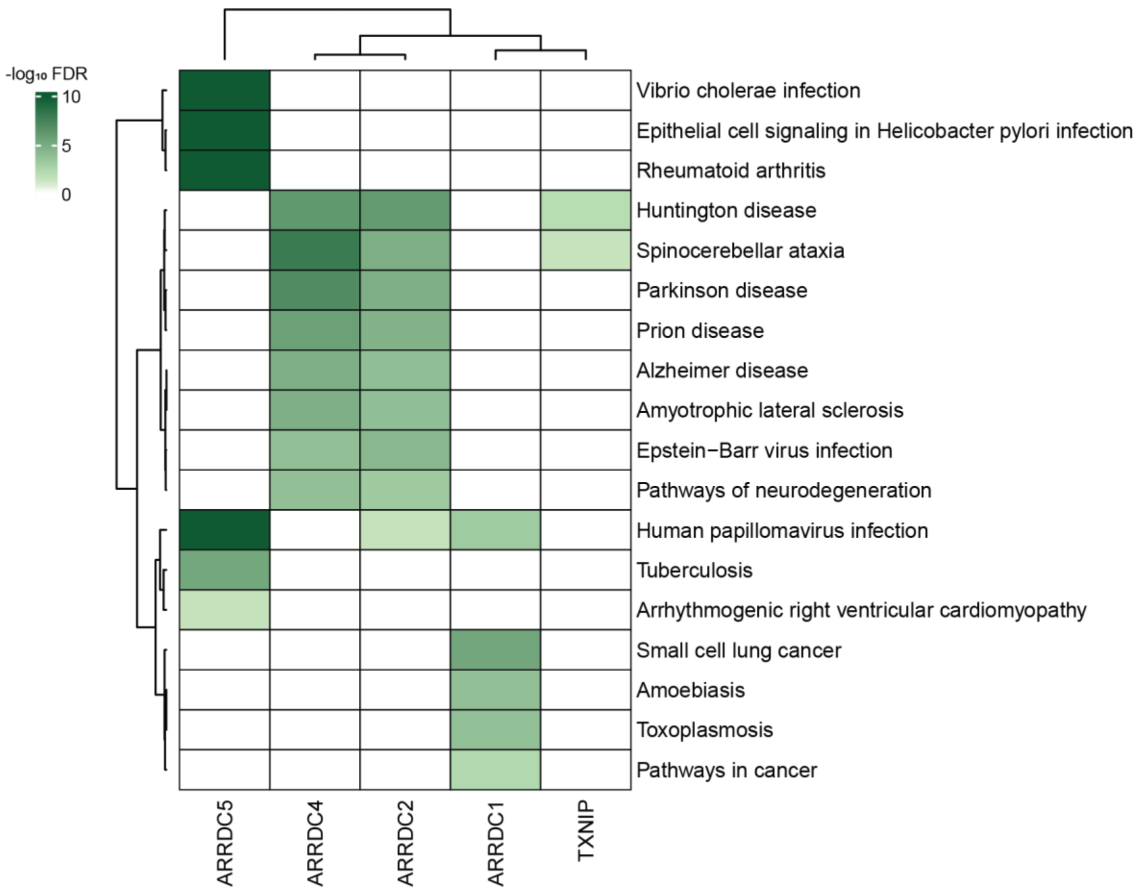


**Figure S9. Association between α-arrestin interactomes and human diseases**

Heatmap depicts disease pathways from Kyoto encyclopedia of Genes and Genomes (KEGG) that are enriched in interactome of each α-arrestin. The significance of the enrichment was tested by enrichR (Kuleshov et al., 2016) and indicated as $-\log_{10}$ FDR. Only the disease pathways that are significantly enriched (FDR < 0.05) are colored.

**Supplementary Table and legends**

**Table S1. List of α-arrestins from human and *Drosophila***

Information about α-arrestin proteins from human **(A)** and *Drosophila* **(B).**

**Table S2. Evaluation sets of α-arrestins PPIs**

Positive and negative PPIs of α-arrestins for human (**A and B**) and *Drosophila* (**C and D**), respectively.

**Table S3. Summary tables of SAINTexpress results**

Summary tables of SAINTexpress results for human **(A)** and *Drosophila* **(B)**.

**Table S4. Protein domains and short linear motifs in the α-arrestin interactomes**

Summary of protein domains and short linear motifs annotated in the interactome of each α-arrestin for human **(A and B)** and *Drosophila* **(D and E)**. Annotated interactions between the short linear motifs in α-arrestins and protein domains in the interactomes from the ELM database are also summarized for human **(C)** and *Drosophila* **(F)**.

**Table S5. Enriched Pfam domains in the α-arrestin interactomes**

Results of enrichment test of Pfam domains in the interactome of each α-arrestin for human **(A)** and *Drosophila* **(B)**.

**Table S6. Subcellular localizations of α-arrestin interactomes**

Summary tables of subcellular localizations of α-arrestins interactomes for human **(A)** and *Drosophila* **(B)**.

**Table S7. Summary of protein complexes and cellular components associated with α-arrestins**

Results of the protein complex enrichment analysis tool (COMPLEAT) and enrichment test of cellular component GO terms for human **(A and B)** and *Drosophila* **(C and D)**.

**Table S8. Orthologous relationship of α-arrestin interactomes between human and *Drosophila***

Predictions of orthologs for α-arrestins and their interacting proteins between human and *Drosophila* using DIOPT **(A and B)**. Results of enrichment test of GO terms (biological process, molecular functions and BP, MF, and KEGG pathway) in the identified orthologs are also summarized for human **(C)** and *Drosophila* **(D)**.

| | ATAC-seq | | | RNA-seq | |
|---|---|---|---|---|---|
| Sample | Properly paired reads (%) | Filtered/dedup reads* | Called peaks | Filtered reads (%) | Alignable reads |
| siCon1 | 117,217,586 (97.8%) | 17,929,628 | 74,373 | 41,756,384 (99.8%) | 37,548,784 |
| siCon2 | 203,055,772 (97.7%) | 31,497,080 | 141,799 | 41,900,786 (99.4%) | 36,139,515 |
| siTXNIP1 | 123,045,656 (97.8%) | 20,050,776 | 69,431 | 41,729,984 (99,8%) | 37,185,131 |
| siTXNIP2 | 179,673,798 (98%) | 25,159,908 | 125,301 | 39,503,312 (99.5%) | 33,418,535 |

**Table S9. Summary of ATAC- and RNA-seq read counts before and after processing**

For ATAC-seq, the number of properly paired reads, filtered/deduplicated reads, and identified narrow peaks are summarized. For RNA-seq, the number of filtered and alignable reads are summarized. *Filtered/dedup reads, filtered/deduplicated reads

**Table S10. Differential accessibility of ACRs and gene expression**

Summary of differential accessibility of ACRs **(A)** and gene expression **(B)** between control and TXNIP depleted condition in HeLa cells.

**Table S11. Summary of ATAC-seq peaks located in promoters and gene expression level**

**(A)** Profiles of ATAC-seq peaks located in promoters of genes. Changes in peak intensities and gene expression levels are also summarized**. (B)** List of genes that exhibited decreased chromatin accessibility at their promoter and decreased RNA expression upon TXNIP knockdown.

| Primer name | Forward/reverse | Sequence | Application |
|---|---|---|---|
| alpha-tubulin | Forward | CTGGACCGCATCTCTGTGTACT | RT-qPCR |
| | Reverse | GCCAAAAGGACCTGAGCGAACA | |
| TXNIP | Forward | GCTCCTCCCTGCTATATGGAT | |
| | Reverse | AGTATAAGTCGGTGGTGGCAT | |
| CD22 | Forward | GCGCAGCTTGTAATAGTTGGTGC | |
| | Reverse | CACATTGGAGGCTGACCGAGTT | |
| L1CAM | Forward | TCGCCCTATGTCCACTACACCT | |
| | Reverse | ATCCACAGGGTTCTTCTCTGGG | |
| CD22 | Forward | GCGCAGCTTGTAATAGTTGGTGC | |
| | Reverse | CACATTGGAGGCTGACCGAGTT- | |
| OTULINL | Forward | GTGTGGAGGCAGAGGTTGAT | |
| | Reverse | ATGCCGCCAAAATAGCTCCT | |
| PRR5L | Forward | GCGGCTGTTGAAGAGTGAAC | |
| | Reverse | AGCCAGAACCTCAATGCGAT | |
| SDC3 | Forward | CTCCTGGACAATGCCATCGACT | |
| | Reverse | TGAGCAGTGTGACCAAGAAGGC | |
| GAPDH | Forward | ATCACCATCTTCCAGGAGCGA | |
| | Reverse | CCTTCTCCATGGTGGTGAAGAC | |
| CD22 #1 | Forward | CGCTGGAGAAGTGAGTTCGG | ChIP-qPCR |
| | Reverse | TCCCTGCCTCCACTGATAGC | |
| CD22 #2 | Forward | GACGCTGAGATGAGGGTTGG | |
| | Reverse | TGACTCAGGAGGTTGGCAGA | |
| CD22 #3 | Forward | TCCCCACTCTTCTCGCTCTC | |
| | Reverse | ATTTGCGAGGTTGAGGTTGTC | |
| L1CAM #1 | Forward | CAGCTCAGTGCCTCATGGAA | |
| | Reverse | GAGACTGCTTCCAGAGTGGG | |
| CD22 #2 | Forward | GGAATGCTTCACTGGGCAAC | |
| | Reverse | GGGGTAAGAATTCCGGAGCC | |
| CD22 #3 | Forward | CGTGTCTGAGAAAGGAAGCCA | |
| | Reverse | CGGCTTATCCCGATCTACCC | |

**Table S12. List of primer sequences used in this study**
